# Estimating sequence diversity of prion protein gene (*PRNP*) in Portuguese populations of two cervid species: red deer and fallow deer

**DOI:** 10.1101/2022.08.16.504133

**Authors:** Jorge C. Pereira, Nuno Gonçalves-Anjo, Leonor Orge, Maria A. Pires, Sara Rocha, Luís Figueira, Ana C. Matos, João Silva, Paula Mendonça, Paulo Carvalho, Paula Tavares, Carla Lima, Anabela Alves, Alexandra Esteves, Maria L. Pinto, Isabel Pires, Adelina Gama, Roberto Sargo, Filipe Silva, Fernanda Seixas, Madalena Vieira-Pinto, Estela Bastos

**Affiliations:** Centre for the Research and Technology of Agro-Environmental and Biological Sciences (CITAB), University of Trás-os-Montes e Alto Douro (UTAD), Vila Real, Portugal; Polytechnic Institute of Castelo Branco (IPCB), Castelo Branco, Portugal; Pathology Laboratory, UEISPSA, National Institute for Agricultural and Veterinary Research (INIAV), I.P., Oeiras and Vairão, Portugal; Animal and Veterinary Research Centre (CECAV), AL 4AnimalS, UTAD, Vila Real, Portugal

**Keywords:** cervid, CWD, prion, PRNP, Portugal, susceptibility

## Abstract

Among the transmissible spongiform encephalopathies (TSEs), chronic wasting disease (CWD) in cervids is now a rising concern in wildlife within Europe, after the detection of the first case in Norway in 2016, in a wild reindeer and until June 2022 a total of 34 cases were described in Norway, Sweden and Finland. The definite diagnosis is *postmortem*, performed in target areas of the brain and lymph nodes. Samples are first screened using a rapid test and, if positive, confirmed by immunohistochemistry and Western immunoblotting. The study of the genetics of the prion protein gene, *PRNP*, has been proved to be a valuable tool for determining the relative susceptibility to TSEs. In the present study, the exon 3 of *PRNP* gene of 143 samples from red deer (*Cervus elaphus*) and fallow deer (*Dama dama*) of Portugal was analyzed. Three single nucleotide polymorphisms (SNPs) were found in red deer – codon A136A, codon T98A, codon Q226E – and no sequence variation was detected in fallow deer. The low genetic diversity found in our samples is compatible with previous studies in Europe. The comparison with results from North America, suggests that the free-ranging deer from our study may present susceptibility to CWD, although lack of experimental data and the necessity of extensive survey are necessary to evaluate these populations.

## 1. Introduction

Transmissible Spongiform Encephalopathy (TSE) or prion diseases are a family of neurodegenerative diseases caused by lethal infectious pathogens called Prions. Prions are proteinaceous infectious particles consisting of PrP^sc^, an abnormally folded and infectious isoform of the endogenous prion protein (PrP^C^). Prions can convert PrP^C^ into PrP^sc^, leading to accumulation of aggregated PrP^sc^ in the central nervous system and ultimately causing the animal’s death [1]. TSEs can affect several mammalians species including Humans [2] but show a predominance in ruminants: scrapie in sheep and goats [3] bovine spongiform encephalopathy (BSE) in bovids [4] and chronic wasting disease (CWD) in cervids [5].

The term ‘wasting disease’ is due to most significative feature of CWD in cervids, the pronounced weight loss [6], but the most important and dangerous trait of CWD is the effortless capacity of transmission between animals, resulting in endemic infections within susceptible populations [7]. Studies in large range of cervids pointed out that PrP^sc^ has been detected in the lymphatic system, salivary gland, intestinal tract, muscles, and blood, as well as urine, saliva, and feces of infected cervids, which imply to remain infectious in environment for a long time (bound to the soil and plants). This led to a possible horizontal transmission between both farmed and free-ranging cervids [8–10].

The first case identified as CWD occurred in 1967, in a captive mule deer in Northern Colorado, USA [11], but rapidly spread to 26 USA states and Canada [12]. In 2016, the first case of CWD appeared in Europe in a free-ranging reindeer in Norway and further cases were later reported in red deer and moose, ultimately reaching other countries like Finland and Sweden [12,13].

Although the origin of CWD is not well known, the fatality, clinical progression and susceptibility/resistance of CWD has been linked to the genetic variation in functional prion protein gene *PRNP* [14–16]. In cervids the *PRNP* is highly conserved, however, several sequence variations of *PRNP* have been identified and characterized as being implicated in prion disease susceptibility/resistance as well as in CWD disease progression [16,17].

The studies with more impact in cervid *PRNP* genetics have been made in USA and Canada, where both natural and experimental infection of deer species has allowed for the identification of *PRNP* sequence variations that are associated with reduced incidence of disease and/or slower disease progression. Since the identification of the first *PRNP* variation in codon 132 (M132L) in wapiti deer suggesting a reduced susceptibility to infection when the 132L variant is present, several other *PRNP* mutations in other cervids have been associated with CWD susceptibility: Q95H, G96S and A116G (white-tailed deer), S225F (mule deer) [16].

The analysis of the *PRNP* diversity in European cervid species is still limited but some studies carried out in Britain, Norway and Sweden, showed that some *PRNP* genetic variations can be related with CWD susceptibility/resistance [18]. The variations S225Y and P242L found in caribou/reindeer were associated to a higher risk of infection and the N138 polymorphism in fallow deer has been associated with natural resistance [12].

In Portugal there are three main species of cervids that belong to the *Cervidae* family: the red deer (*Cervus elaphus*), the roe deer (*Capreolus capreolus*) and the fallow deer (*Dama dama*) [19]. The red and fallow deer are native species from Portugal, and they were re-introduced in several areas along Portugal. No cases of CWD have been reported yet in Portuguese cervids. Nevertheless, the most recent accession of this disease in Portuguese cervid populations was done between 2006 and 2010 as part of a limited survey carried out in the European Union (EU) [20].

The recent identification of several cases of CWD in Scandinavia rang the alarm of authorities regarding the dissemination of this TSE. CWD is endemic and difficult to eradicate putting in high risk the cervid populations. Genetics studies, to evaluate the variation of *PRNP* gene in European cervid populations are of extreme importance in order to estimate the potential susceptibility of populations to emerging CWD and informing risk assessment and control/surveillance strategies. The extent of *PRNP* sequence variation in Portuguese cervids species has not been previously studied, so the main aim of our research, integrated in a project, was the study of protein-coding sequence *PRNP* gene variability in two species of cervids (red deer and fallow deer) in different populations of Portugal in order to evaluate the genetic variability, resistance or susceptibility to CWD, of these cervid species when compared with *PRNP* genotypes found in Europe.

## 2. Materials and Methods

### 2.1. Animal Samples and DNA extraction

This study includes 143 animals of two different species of wild ungulates: red deer (*Cervus elaphus*) and fallow deer (*Dama dama*). All of them were free-ranging animals from different places from east center of Portugal and were hunted or found dead in reserves or hunting grounds. Red deer (n=57) and Fallow deer (n=86) samples come from three hunting/reserves areas: Vale Feitoso [VF] (RD=19; FD=61), Rosmaninhal [RM] (RD=27; FD20) and Termas de Monfortinho [TM] (RD=9; FD=5) (Figure 1).

**Figura 1.**
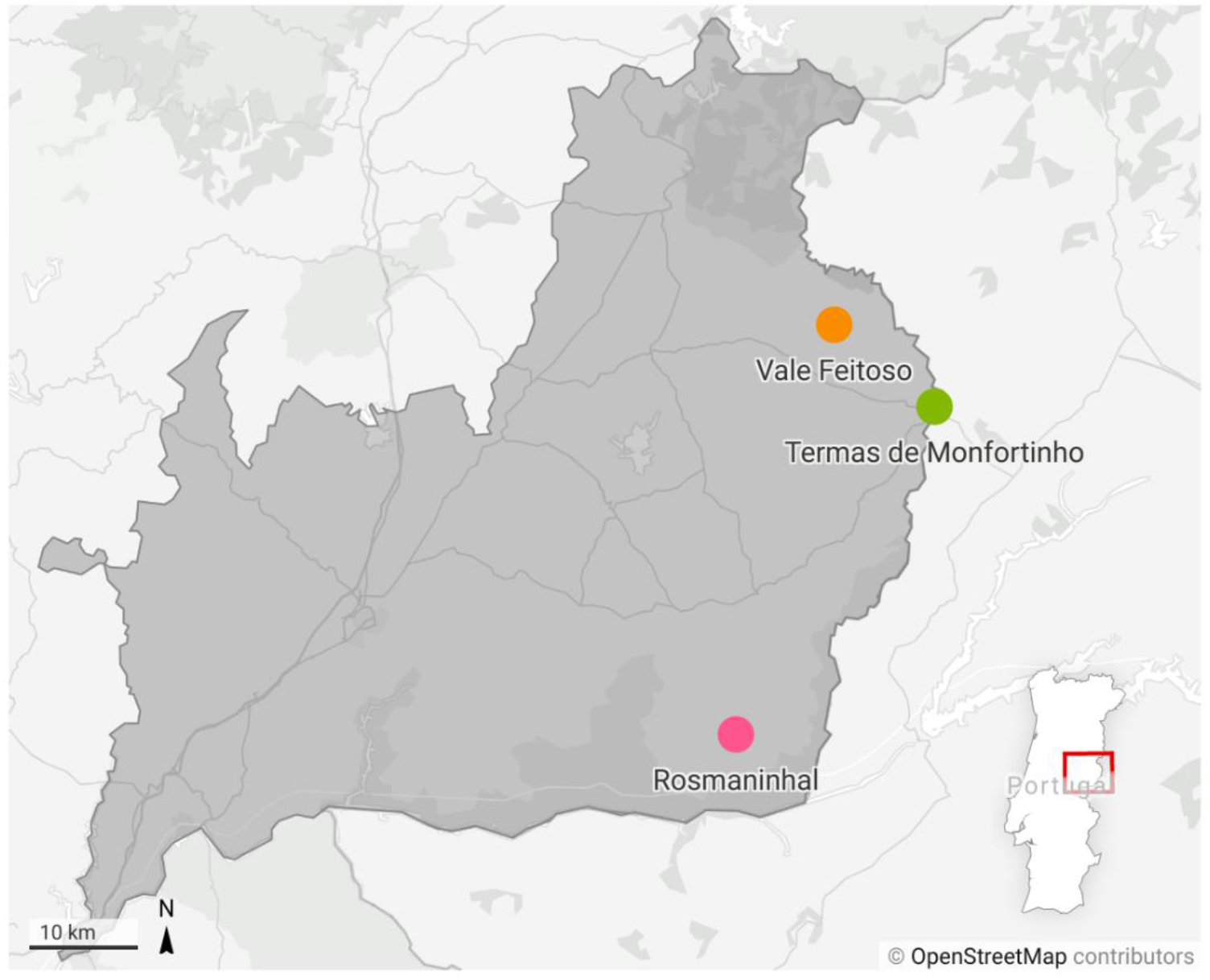
Map of the east center of Portugal (Beira Baixa district) showing the geographic locations where different samples were obtained.

Masseter muscle and retropharyngeal lymph node of each animal were collected for genetic analysis of prion protein gene, *PRNP*.

Genomic DNA was extracted from 50 mg of frozen muscle or lymph node using a NZY Tissue gDNA Isolation kit (NZYTech, Lda. - Genes and Enzymes) following the manufacturers protocol and stored at -20º C for further analysis.

### 2.2. PRNP gene amplification, Sequencing and Statistical analysis

The full exon 3 of *PRNP* gene (771 bp) of all animals was amplified by PCR using MyTaq™ Red Mix (*Bioline Meridian Life Sciences-BIO-25043*). Briefly, in 20 μl reaction: 10 μL MyTaq™ Red Mix, 1 μL of 20 μM primers (F223-ACACCCTCTTTATTTTGCAG and R224-AGAAGATAATGAAAACAGGAAG), 1.75 μL PCR water, 1 μL of 100 ng/μL DNA template. All reactions were carried out using the following protocol: 95°C for 15 minutes, 35x cycles (95°C for 30 seconds, 60°C for 90 seconds, and 72°C for 60 seconds), with a final extension at 72°C for 10 minutes.

PCR products were purified and afterwards were sequenced by Eurofins Genomics GmbH in Germany. Raw sequence reads were proofed, and chromatograms were visually inspected to verify all base changes using SnapGene Viewer v. 5.1.5 together with Unipro UGENE v. 40.0 [21]. Nucleotide sequences were aligned and trimmed to coding region of *PRNP* gene with 771 bp, using Unipro UGENE v. 40.0 [21], by referencing them with sequence of red deer (accession number KT_845862) and fallow deer (accession number MK_103017) obtained from GenBank. Furthermore, these DNA sequences were translated into protein using standard genetic code with the same bioinformatic software. The editing and visualization of protein multiple alignments for the Figure 1 was made in Jalview 2.11.1.4 [22].

Four sequences were summitted to GenBank with the following accession numbers: OP179612, OP179613, OP179614 and OP179615.

### 2.3. PrP^sc^ detection

Previously, all cervids were screened for PrP^sc^ in both medulla oblongata at the level of the obex (350±40 mg) and retropharyngeal lymph node (200 ± 20 mg of tissue from outer cortex) using TeSeE™ SAP Combi Kit Bio-Rad (reference 3551191) for purification and ELISA detection of PrP^sc^ following manufacturer’s instructions.

## 3. Results

All cervids screened in the present study resulted negative for PrP^sc^ test.

The comparison of the coding region of *PRNP* gene (771 bp) and the corresponding protein sequence - PrP^C^ (256 aa) from 57 red deer samples collected in Portugal, showed high conservation. Nevertheless, three sequence variations from both consensus sequences were identified: one silent or synonymous variation at position 408 (codon 136) (gcT/gcC), and two non-synonymous variations at positions 292 (Acc/Gcc), and 676 (Cag/Gag), resulting in amino acid changes in codons 98 (substitution of threonine (T) with alanine (A)) and 226 (substitution of glutamine (Q) to glutamic acid (E)) (Table 1). The synonymous mutation at position 408 (codon 136) showed to be linked to the non-synonymous mutations at position 676 (codon 226), producing haplotypes: refer as Q226(t408-c676) or as E226(c408-g676).

**Table 1.**
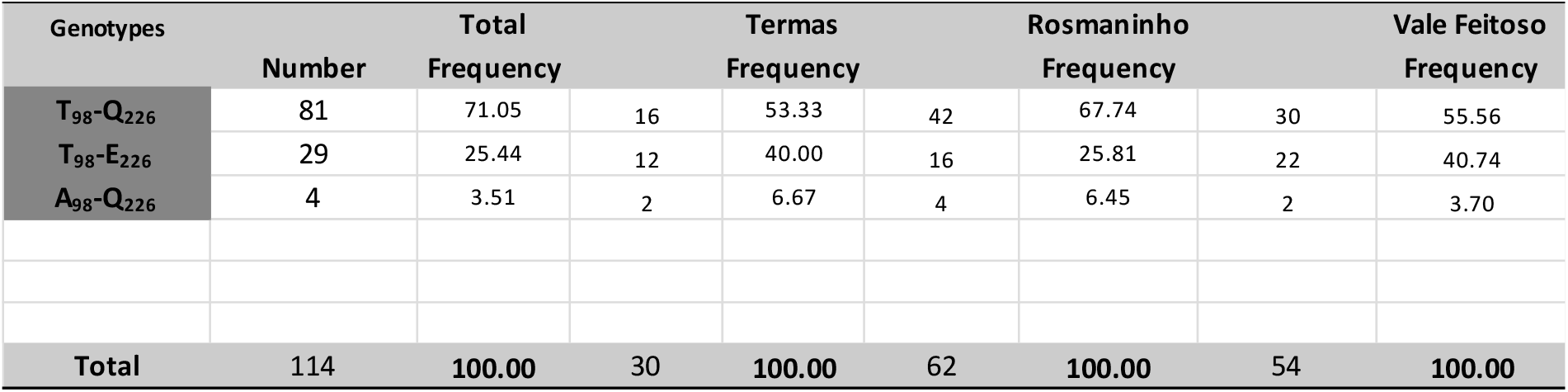
Amino acid variation within the exon 3 and haplotype frequencies of cervid *PRNP* in red deer populations.

The genotypic and haplotype frequencies were calculated for all samples collected, and separately for the different populations (Tables 1 and 2).

**Table 2.**
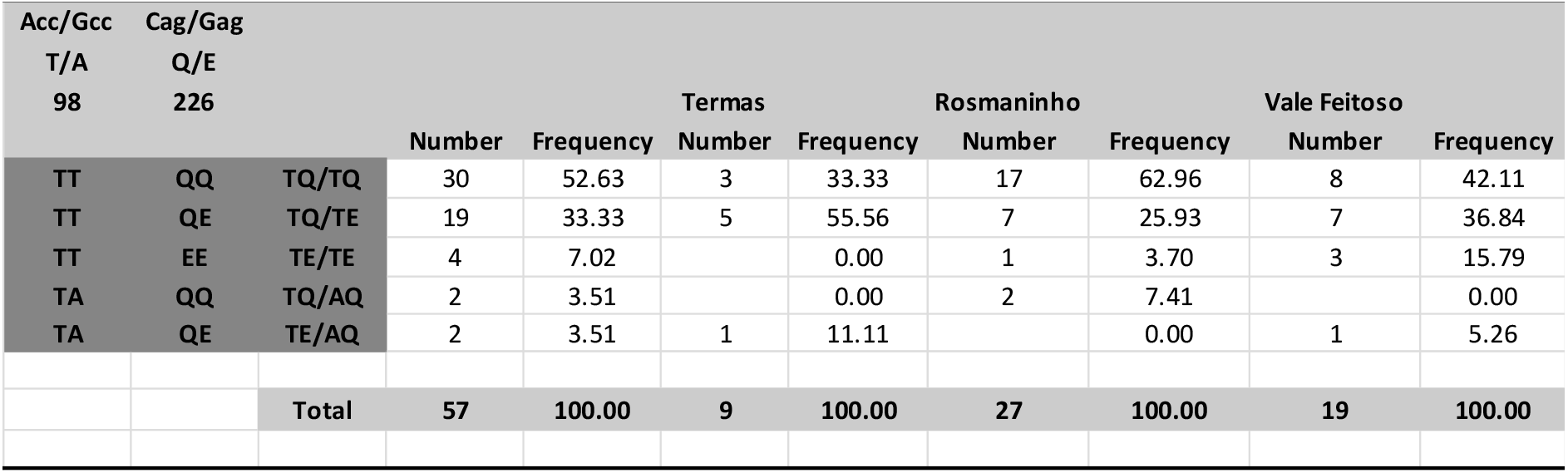
Genotype frequencies of *PRNP* polymorphisms in Portuguese red deer populations.

Observing the table 1, the non-synonymous variations produced three haplotypes in the Portuguese red deer populations: T_98_-Q_226_ (TQ), T_98_-E_226_ (TE) and A_98_-Q_226_ (AQ) with haplotype frequencies of 71%, 25.4% and 3.5%, respectively. These haplotypes are present in all different geographic populations evaluated with different frequencies, being the haplotype TQ the one with highest frequencies (67%-53%), and the AQ haplotype the one with lowest frequencies, ranging from 6.7%-3.7%.

The *PRNP* coding region analysed in 77 fallow deer samples showed no sequence variability in the population. All the PrP^C^ amino acid sequence from samples corresponded to the red deer haplotype TE with the exception of a single amino acid position (codon 138) that contains asparagine (N) in fallow deer and serine (S) in red deer (Figure 2).

**Figure 2.**
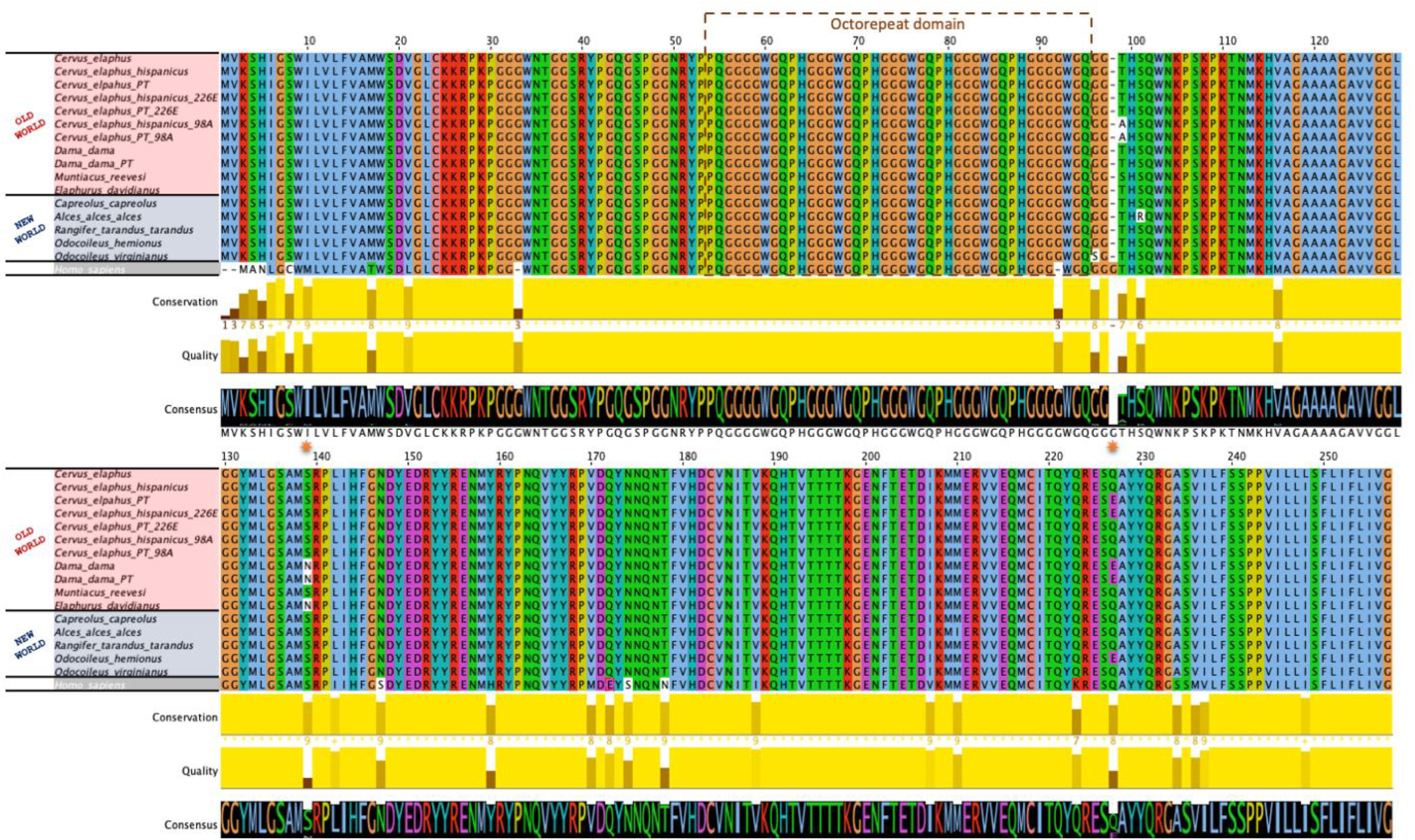
Protein sequence alignment of New and Old World cervid showing conserved homology between species and other molecular conserved features.

Another molecular feature found in the red deer and fallow deer samples and usually associated with consensus amino acid sequence, is the amino acid makeup of the terminal signals and the octorepeat domain (5 peptide repeats with 3 octapeptides of PHGGGWGQ flanked by 2 nonapeptides of P[Q/H]GGGGWGQ (Figure 2).

## 4. Discussion

This study represents the first survey of *PRNP* genetic variation in Portuguese wild (or free-ranging) cervid populations. A total of 143 individuals were included in the present study: 57 red deer and 86 fallow deer, distributed in three collection areas.

The *PRNP* coding region, corresponding to exon 3, is extremely conserved among cervid species, and in some cases, like the fallow deer, there is no intra species variation. However, the main goal of sequencing this *PRNP* region is to identify possible DNA sequence variations that could lead to important amino acid substitutions in PrP^C^ sequence previously associated with lower rates and/or delayed CWD disease progression [12,18].

Many studies have presented several non-synonymous polymorphisms in PRNP protein sequence, but there is still no data that can relate any amino acid variation in European red deer with susceptibility to CWD [18]. The analysis of *PRNP* gene and protein sequence in Portuguese red deer revealed only two amino acid substitutions - T98A and Q226E – but none of the amino acid substitutions was related to CWD susceptibility/resistance. Other non-synonymous polymorphisms already identified in red deer like the V15A, G59S, M208I were not observed in red deer samples.

Regarding the first amino acid substitution identified in this study, T98A, although it is not directly related with CWD susceptibility, some studies question/highlighted the proximity of this variation in red deer with the variation in codons 95 and 96 found in the white tail deer, which was proved to be related with reduced CWD susceptibility [23,24]. Besides, studies of CWD on camels revealed the natural development of prion disease when Alanine is present on this location [25].

The mutation on codon Q226E is of great importance since it is located in α-helix 3, where part of the protein is involved in pathogenic processes during prion disease. In fact, amino acids variations can influence protein stability and cause different conformation of PrP^C^ protein [26]. The Portuguese red deer samples present homozygotic - Q and E - and heterozygous genotypes. Studies in transgenic mice expressing cervid PrP with Q226E variation referred that it may contribute to differences in disease progression and susceptibility to certain prion strains [27], but in other experimental assays of oral transmission of CWD using deer with different genotypes, E226E, Q226Q and Q226E, showed that all animals became infected [28]. Another interesting fact is that the presence of polymorphism S225F in mule deer [29] is associated with reduced CWD susceptibility but in Caribou/Reindeer S225Y variation is related with higher risk of infection [15]. There is still no main conclusion of homozygous or heterozygous or adjacent amino acid variations and their relationship with CWD susceptibility, due mainly to the lack of concrete and specific studies on this SNPs.

From the analysis of *PRNP* gene and protein sequence in Portuguese red deer three different haplotypes were identified T_98_-Q_226_ (TQ), T_98_-E_226_ (TE) and A_98_-Q_226_ (AQ). These haplotypes are not new and have been previously described in Italian, Scottish [26], Spanish [30], British, Norwegian and Czech red deer populations [24]. The haplotypes TQ and TE revealed a high haplotypic frequency in all the samples and geographic regions studied. When compared with other European red deer populations it is possible to observe similar frequency values with Spanish, Italian and Czech populations. Regarding the AQ haplotype, the total haplotypic frequency is lower than the ones found in Spanish, Italian and Czech populations, but similar to the Scottish samples. It is possible that the lower number of analyzed animals could influence this frequency values in our population. The high frequency of TQ haplotype found in our study, is identical to *PRNP* sequences found in North American cervids that are highly susceptible to CWD. Moreover, the red deer samples analyzed in this study were homozygous in codon M132 and G96.

As already described in several publication, the mutation of codon M132L has been associated with some degree of protection against CWD, as well as with delaying in its progression. Mutation in codon G96S has been associated with reduced susceptibility to CWD [31], which means homozygous genotypes 96G could have high susceptibility to CWD. The high frequency, together with the homozygous state of codons M132 and G96 and the fact that the same condition has been observed in European red deer populations, suggests that the analyzed populations might be at high risk to be very susceptible to CWD [24].

The analysis of fallow deer samples showed no intra species variation in exon 3 of *PRNP* gene and consequently in PrP^C^ protein sequence. This monomorphic state was previously found in other European fallow deer populations, as well as in other species such as the roe deer and the axis deer. The haplotype TE with N138 of fallow deer was observed in all samples, which is interesting since the codon 138 with asparagine (N) has been linked to resistance to natural infection and prolonged incubation periods in intra-cerebrally infected animals, following previous experimental studies [12,18].

The sequence variation in the coding region of *PRNP* as an important genetic agent that can influence the expression of TSEs, but there are other genetic variations in the *PRNP* gene that could be related with the expression of TSEs - indels in the promoter region, frequency of octarepeats -, thereafter possible CWD in cervids [32].

The variation in the number of octarepeat units (PHGGGWGQ) in the PRNP gene is another genetic marker that can be used to study the susceptibility/resistance of cervids to CWD, although no studies were made in cervids, there are studies in scrapie, BSE and CJD [33–35]. In our study, in both species, the octorepeat domain as represented by 5 peptide repeats with 3 octapeptides of PHGGGWGQ flanked by 2 nonapeptides of P(Q/H)GGGGWGQ) without any polymorphism, same as found in Vázquez-Miranda and Zink [36] and Zink [32]. When compare with human octorepaeat domain only a single polymorphism is found (position 92). We think it is important to the future study of CWD in cervids, the analysis of the relationship between PRNP genotype, indels in the promoter region and frequency of octarepeats.

In summary, this survey has provided an analysis of sequence variation of *PRNP* exon 3 and consequently of prion protein sequence - PrP^C^ - in two Portuguese free ranging cervid species, the red deer and the fallow deer. The results were compared with European populations and as well with North American cervids affected by CWD. All animals in the present study, were previously screened and were negative for the presence of CWD. The *PRNP* gene and PrP^C^ analysis showed a high degree of conservation among and between the two species as expected from other studies. Although the effect of the haplotypes found in our study (and that are also present in other European species) on CWD susceptibility are not determined, some of the identified SNPs indicate that our populations might be susceptible to CWD.

CWD is a high contagious TSEs that is difficult to eradicate, with a quick expansion among states, countries and continents, and affecting not only farm but also wild cervids populations. The first cervid found with CWD infection in Europe (Norway) in 2016 was probably the turnover point for CWD studies, not only because there is no certainty of the origin of CWD but also because at that time complete strategies against CWD dissemination were inexistent. In our opinion, multi-disciplinary approaches including genotyping, PrPres detection, identification of risk factors and others, are of great importance to evaluate the risk of occurrence of CWD in Europe and more specific in Iberian Peninsula.

## Author Contributions

All authors have read and agreed to the published version of the manuscript.

